# Testing the unitary theory of language lateralisation using functional transcranial Doppler sonography in adults

**DOI:** 10.1101/437939

**Authors:** ZVJ Woodhead, AR Bradshaw, AC Wilson, PA Thompson, DVM Bishop

## Abstract

Cerebral lateralisation for language can vary from task to task, but it is unclear if this reflects error of measurement or independent lateralisation of different language systems. We used functional transcranial Doppler sonography to assess language lateralisation in 37 adults (7 left-handers) on six tasks, each given on two occasions. Tasks taxed different aspects of language function. A preregistered structural equation analysis was used to compare models of means and covariances. For most people, a single lateralised factor explained most of the covariance between tasks. A minority, however, showed dissociation of asymmetry, giving a second factor. This was mostly derived from a receptive task, which was highly reliable but not lateralised. The results suggest that variation in strength of language lateralisation reflects true individual differences and not just error of measurement. Inclusion of several tasks in a laterality battery makes it easier to detect cases of atypical asymmetry.

## Introduction

Hemispheric dominance for language is often assumed to be unidimensional and consistent across language domains, but this assumption can be questioned (Bishop, 2013; Bradshaw, Thompson, Wilson, Bishop, & Woodhead, 2017). Discrepant laterality across different language tasks (e.g. Gaillard et al., 2004; Stroobant, Buijs, & Vingerhoets, 2009; Tailby, Abbott, & Jackson, 2017) could be simply due to measurement error (Ramsey, Sommer, Rutten, & Kahn, 2001); alternatively, task differences may represent meaningful individual variation in the hemispheric organization of different language networks. It has been difficult to distinguish these possibilities, because, while we have ample evidence that the left hemisphere is heavily implicated in language function at the group level, relatively little is known about the reliability of lateralization in individuals. It is evident that a standard model based on average brain activation may give a misleading impression of uniformity (Seghier & Price, 2018). Furthermore, there is evidence that there may be subgroups of people with distinct laterality profiles, related to handedness (Mazoyer et al., 2014). Such variability in cerebral lateralisation may have functional significance, for example in terms of impaired language abilities (Bishop, 2013). In clinical neurosurgical contexts, it is important to know whether a single indicator of an individual’s language laterality is sufficient, or whether a battery of measures is needed to capture laterality in multiple language domains (Gaillard et al., 2004; Stroobant et al., 2009; Tailby et al., 2017). Before we can make headway in answering such questions, we need to have reliable measures.

Here we report a study using functional transcranial Doppler sonography (fTCD; Knecht et al., 1998) to measure speed of blood flow in left and right middle cerebral arteries (a proxy for neural activity in language-related areas of the brain) during six different language tasks (tasks A-F). The fTCD data were used to derive laterality indices (LIs), which quantify the balance of activation in left and right hemispheres. All participants were tested on the whole battery in two separate sessions on different days in order to estimate the reliability of the LIs and the extent to which lateralization of different tasks could be explained in terms of a common factor.

### Laterality at the level of the population and the individual

The question of whether language lateralisation is a unitary function has two distinct interpretations: (a) whether there are differences in extent of lateralisation across different language functions or (b) whether there are individual differences in how the strength of lateralisation varies across language functions. We first review existing literature on these questions and then present simulated data to show how predictions made by the two accounts are independent and additive, but can be tested within a common framework (structural equation modelling, SEM).

### Task-related variation in extent of language lateralisation

Most theories of language lateralisation have focused on how language functions are lateralised in the brain in typical humans. Such theories are not concerned with individual differences, but make theoretical statements about the properties of language that are associated with lateralised activity. An influential example of such a theory is Hickok and Poeppel’s dual route model of speech processing (Hickok & Poeppel, 2007). This contrasts a dorsal stream from superior temporal to premotor cortices via the arcuate fasciculus, which is associated with sensorimotor integration of auditory speech sounds and articulatory motor actions; and a ventral stream from temporal cortex to anterior inferior frontal gyrus, which is involved in access to conceptual memory and mapping of sound to meaning (Rauschecker, 2018). Hickok and Poeppel proposed that the dorsal stream is left lateralized, whereas the ventral stream is bilateral. This kind of theory makes predictions about task-related differences that can be assessed by comparing mean LIs in a sample. Thus, the prediction from the dual route model is that mean LIs for tasks involving the dorsal stream will show left-lateralisation, whereas LIs from tasks primarily involving the ventral stream will not be lateralised.

Hickok and Poeppel’s model contrasts with other theoretical accounts. For instance, Dhanjal et al proposed that left lateralization was a characteristic of tasks involving lexical retrieval (Dhanjal, Handunnetthi, Patel, & Wise, 2008). Evidence came from an fMRI study investigating propositional speech (e.g. sentence generation) and non-propositional speech (e.g. reciting memorized speech): articulatory jaw and tongue movements and non-propositional speech co-activated bilateral dorsal areas, including the superior temporal planes, motor and premotor cortices. Only the lexical retrieval component of propositional speech resulted in left lateralized activity (in the inferior frontal gyrus and premotor cortex).

Yet other accounts have focused on the complexity of the speech stimulus (Peelle, 2012), or argued that lateralization is specifically linked to aspects of complex syntactic processing (Bozic, Tyler, Ives, Randall, & Marslen-Wilson, 2010; Friederici, 2011).

### Individual differences in cerebral lateralisation

Discussions about the nature of language lateralization are complicated by individual differences; although most people show the typical pattern of language laterality, some individuals show the reverse pattern – right-hemisphere language. In a large-scale comparison of left-and right-handers, Mazoyer et al (2014) reported that strong right-hemisphere bias for a sentence generation task was seen exclusively in left-handers, though milder departures from left hemisphere dominance were seen in right-as well as left-handers. A subset of people with bilateral language has also been described for many years (Milner, Branch, & Rasmussen, 1966), but this category is ambiguous. These could be people who engage both hemispheres equally during language tasks, or people who are strongly lateralized for different tasks, but in different directions. This latter scenario would provide strong evidence against a unitary hypothesis, by demonstrating that a person’s language laterality could not be predicted by a single dimension.

Individual differences in cerebral lateralisation have previously been observed in the comparison between left lateralised verbal functions versus right lateralised nonverbal functions. This might suggest complementarity of the two functions within the brain; however, where individual differences in these biases have been assessed, several studies have found them to be dissociated (Badzakova-Trajkov, Corballis, & Häberling, 2016; Groen, Whitehouse, Badcock, & Bishop, 2012; Rosch, Bishop, & Badcock, 2012; Whitehouse & Bishop, 2009; Zago et al., 2015; cf: Cai, Van der Haegen, & Brysbaert, 2013; Vingerhoets et al., 2013). Again, handedness has been noted as an important factor, with right-handers showing less evidence of complementarity of verbal and visuospatial functions than left-handers (Zago et al., 2015). Here, we consider whether similar dissociations might be found *within* the domain of language. Although previous investigators have considered association or dissociation in average patterns of activation for different tasks (Hesling, Labache, Jobard, & Leroux, 2018; Pinel & Dehaene, 2010), there has been little previous research documenting individual differences in task-related variation. Inconsistent LIs from task to task could simply reflect noisy measurement, making dissociations hard to interpret. Thus, in order to throw light on individual differences in language laterality, we need to include repeated measures, so that reliability of LIs from different tasks can be assessed.

### Simulated data to illustrate predictions

It is possible to integrate models of task variation in lateralisation with a model of individual differences in the kind of framework shown in Figure 1. For simplicity, this shows simulated data on just two tasks, A and B, to contrast predictions from different models of the structure of language lateralisation. The Population Bias model is the simplest: it shows a population bias to left-sided language laterality (i.e. positive LI values) that does not depend on the task. There are no consistent individual differences: any variation in laterality is just caused by random error. This is not a very plausible model, but provides a useful starting point from which to build more complex scenarios. Formally, the function for predicting an individual’s LI is as follows:

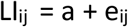

where *i* indexes the task, and *j* the individual, *a* is an intercept term corresponding to population bias, and *e* is random error.

In the Population Bias model, the mean LIs for different language tasks (shown by the horizontal and vertical red dotted lines) are all the same and equal to *a* (in this case set to 1). Note that because there are no stable individual differences, the correlations between LIs for the same task measured on different occasions (left hand panel), and between different tasks measured on the same occasion (right hand panel) are zero.

The second model is the Task Effect model. This incorporates consistent task-specific variation, without any stable individual differences. Formally,

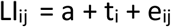

where t_i_ is a task-specific term. The only difference from the Population Bias model is that the means differ for different tasks – i.e. tasks A and B have mean LIs of 1 and 2 respectively. Again, variation in individuals’ LI scores is due to random error (e), rather than any systematic individual differences, as evidenced by zero test-retest correlations.

The next model is a Person Effect model. This includes stable individual differences: a person’s score on any test occasion depends on an intrinsic lateral bias, which is constant from task to task but varies from person to person, i.e.

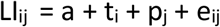

where p_j_ is the person-specific term. This model predicts significant correlations between the same task tested on different occasions, and different tasks tested on the same occasion. An important point is that these correlations depend solely on the relative contribution of individual difference (p) vs random noise (e) to the LI. It does not matter whether there are also task-related effects (t) on the LI. Thus, in the example, we have one task that is lateralised (mean LI of 2) and one that is not (mean LI of 0), yet on this model, the test-retest correlation for either task will be the same, and equivalent to the cross-task correlation.

The final model incorporates a Task by Person Effect: i.e., there are stable individual differences that show up as significant test-retest reliability on any one task, but the rank ordering of lateralisation varies from task to task, so cross-task correlations are low. Formally:

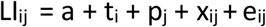

where x_ij_ reflects a contribution that is specific to the task and the individual. The depicted scenario in Figure 1 is an extreme one, with no relationship between a person’s laterality on tasks A and B; in practice, there could be significant cross-task correlations, but if the within-task correlations are higher than cross-task correlations, then this would be evidence that individual differences in laterality are to some extent task-specific.

A key point illustrated by these simulations is that testing the multivariate model of language laterality at the population level requires different evidence – i.e. testing between means – than a multivariate model of individual differences, which requires us to consider correlations within and between tasks. Furthermore, predictions from these two types of model are independent, because correlations are not influenced by mean values. We can use structural equation modelling (SEM) to evaluate the relative fit of these four models to data on language lateralisation for participants who have LIs assessed on a range of tasks on two occasions.

**Figure 1.**
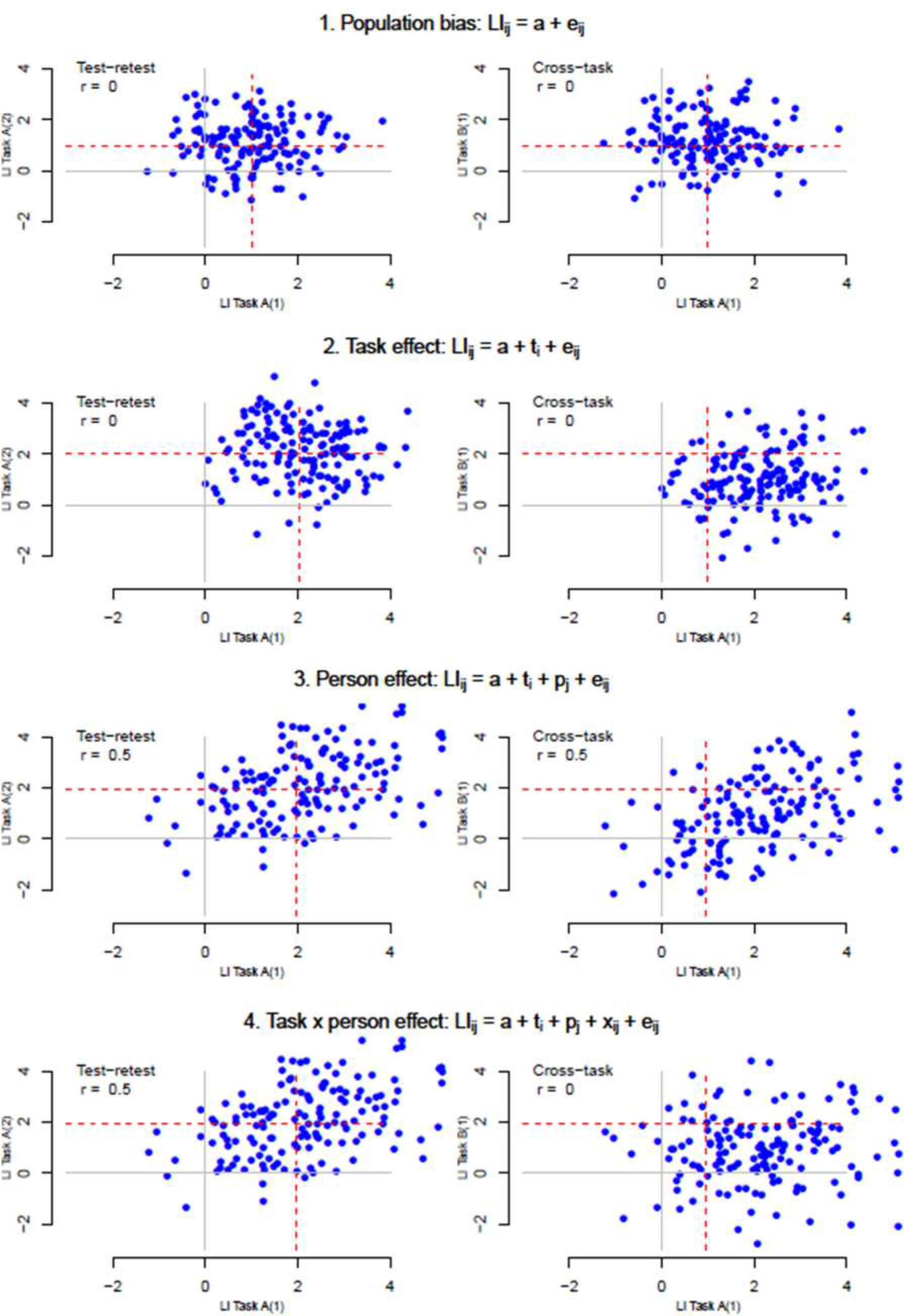
Simulated data of different theoretical models of variance across sessions (1 and 2) and tasks (A and B) in language lateralization. Red dotted lines show the mean lateralization index (LI) for the task / session.

### Hypotheses

We preregistered a set of hypotheses that were tested through SEM model comparison, as described in the Methods below.

We first tested two hypotheses concerning the group mean LI values. First, we tested the dorsal stream hypothesis (Hickok and Poeppel, 2007), which predicts that strength of lateralization depends on the extent to which tasks map on to the dorsal versus ventral speech processing streams (dorsal = stronger left lateralization). Second, following Dhanjal et al (2008), we tested the lexical retrieval hypothesis, which maintains that lateralization depends on the extent to which tasks require lexical retrieval (more lexical retrieval = stronger left lateralization).

A second set of hypotheses concerned individual differences in LI value. We predicted that a Task by Person Effect model, whereby covariances between tasks were modelled by two latent factors, would give a better fit to the data than a Person Effect model, where covariances were modelled by only one factor.

## Methods

### Preregistration

This project was preregistered on Open Science Framework prior to data collection (https://osf.io/tkpm2/). A number of changes were made to the analysis plan after collection of the data – an updated protocol is documented here: https://osf.io/bjsv8/. Departures to the original protocol are explained in the **Departures from pre-registered methods** section below.

### Design

A test-retest, within-subject design was used. Lateralisation of brain activity was measured using Functional Transcranial Doppler Sonography (fTCD) during six language tasks: (A) List Generation, Phonological Decision, (C) Semantic Decision, (D) Sentence Generation, (E) Sentence Comprehension, and (F) Syntactic Decision. Participants were tested on two sessions spaced by between 3 days and 6 weeks. Hence, each participant provided data from six tasks tested twice (A1-F1, A2-F2).

### Participants

A sample size of n=30 was determined by simulations of data from six tasks administered on two occasions, to determine the smallest sample size that would reliably distinguish data generated from a two factor vs single factor model, and give acceptable fit indices (see laterality_simulations files, https://osf.io/tkpm2/). The simulations were based on the models of covariances, as the factor structure of the measures is our primary interest, and this gave a more conservative power estimate. We note that the sample size is small relative to those usually recruited for SEM analyses. However, because all measures were taken twice, with no practice effects expected (on the basis of previous studies with this method), there are several estimates of most parameters. For instance, the correlation between LIs for tasks A and B is estimated from A1B1, A1B2 and A2B2. Thus the repeated measures give low degrees of freedom relative to the number of measures.

In our original study pre-registration we did not plan to select participants according to handedness. However, both prior literature and our own preliminary data indicated that it would be advisable to treat right-and left-handers separately, as the pattern of associations between language tasks appeared to differ according to handedness, so combining handedness groups could give a misleading picture. We became concerned that results from our pre-registered analysis on 30 participants (7 left-handers) were potentially misleading, as the factor structure that emerged seemed driven by a few left-handers. We therefore tested additional participants to give a total sample of 30 right-handers and seven left-handers, and we report analysis based on this larger sample as exploratory results.

All participants gave written informed consent. Procedures were approved by the University of Oxford’s Medical Sciences Interdivisional Research Ethics Committee (approval number R40410/RE004). Subjects were recruited using the Oxford Psychology Research Participant Recruitment Scheme (https://opr.sona-systems.com) and by poster advertisements. The inclusion criteria were: aged 18-45 years; English native language speakers; and with normal or corrected to normal hearing and vision. Exclusion criteria were: a history of significant neurological disease or head injury; or a history of developmental language disorder.

It was not possible to record a Doppler signal via the temporal window in three participants. In these cases the participant was reimbursed but not tested further, and another participant was recruited in their place. One participant had excessive motion artifacts in his first session, so another participant was recruited in his place. The initial group of 30 participants (17 female and 7 left-handed) had a mean age of 26.0 years (SD = 7.2 years; range: 19.2 to 45.1 years). The final group, including seven additional right-handers (2 females) had mean age 25.9 years (SD = 6.8 years) with the same age range.

### Procedure

The order of the six language tasks was counterbalanced between subject and session. At each session, fifteen trials of each task type were conducted with breaks in between tasks.

### Language tasks

The six tasks were designed to be matched in trial structure, as far as feasible, so that differences in laterality should reflect as far as possible the linguistic task demands. The first five tasks had a visual stimulus on each trial presented against a grey background, to keep the visual demands as similar as possible; the sixth task involved presentation of written words. All stimulus materials are available on Open Science Framework (https://osf.io/8s7vn/).

The rest period prior to stimulus presentation was used for baseline correction to equate the left and right channels. Trials were 33 seconds long, and followed the structure shown in Figure 2. Trials started with the word ‘CLEAR’ on screen for 3 seconds, indicating that participants must clear their mind in preparation for the next trial. The language task followed, lasting for 20 seconds. Procedures for each task type are detailed below, and examples of stimuli are shown in Figure 3. Note that for tasks B, C, E and F, participants made responses to a series of stimuli on each trial to ensure the participant was engaged in language processing throughout the activation interval. Rapid presentation of multiple stimuli in a trial has been shown by Payne et al (Payne, Gutierrez-Sigut, Subik, Woll, & MacSweeney, 2015) to maximise lateralised activation in fTCD. After the task, ‘REST’ appeared on screen for 10 seconds, during which participants were required to clear their minds.

**Figure 2.**
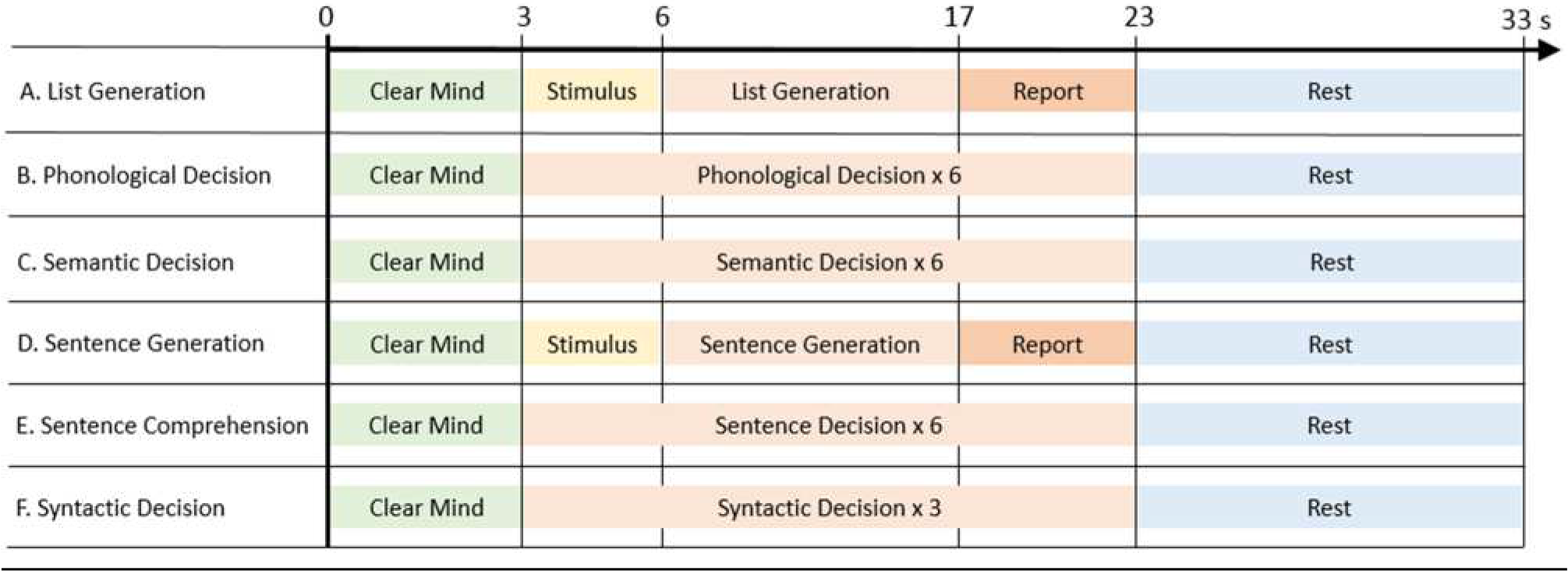
Timings within a single trial for all six task types.

**Figure 3.**
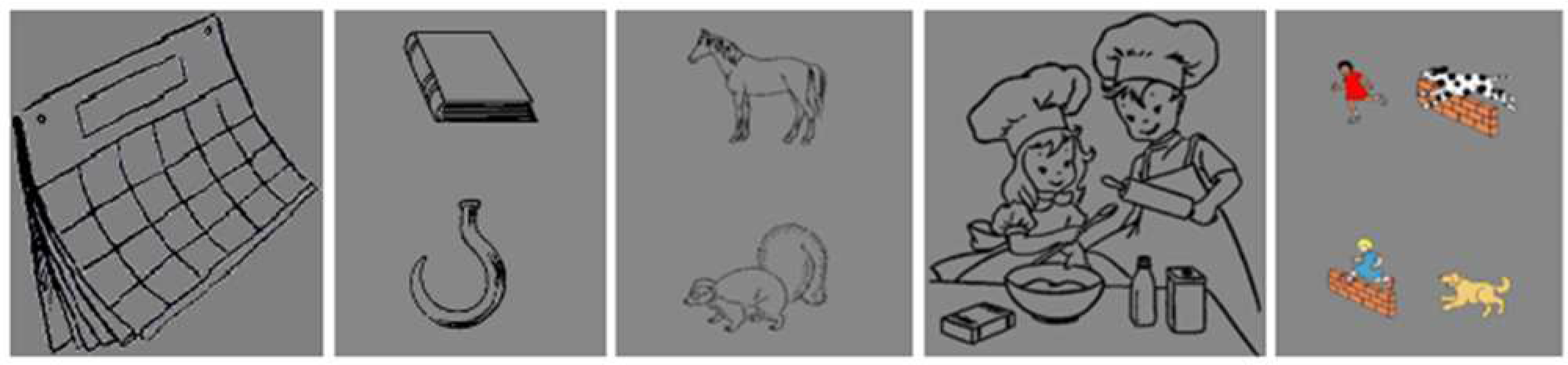
Example stimuli for the language tasks. From left to right: picture stimulus for List Generation task (A; recite months of the year); a matching picture pair (‘book’ / ‘hook’) for the Phonological Decision task (B); a matching picture pair for the Semantic Decision task (C); picture stimuli for the Sentence Generation task (D); and a picture pair for the Sentence Comprehension task (E; ‘The dog chases the girl who is jumping’).

#### A. List Generation

This task was based on the reference task used by Mazoyer et al (2014). Participants were asked to recite an automatic sequence of words (non-propositional speech) in response to a picture. In each trial, a line drawing was displayed on a grey background for 3 seconds. Participants were trained to produce different sequences for different pictures: reciting the numbers from 1-10, the letters from A-J, the days of the week or the months of the year. A fixation cross was then presented in the center of the screen for 11 seconds, during which the participant recited the words covertly (silently) in their head. Following this, a ‘REPORT’ prompt was shown for 6 seconds, indicating that participants should say the sequence aloud. The list generation task involves generation of phonological output, and so should index the dorsal stream, but because it involves repeated, overlearned material, it does not implicate the ventral stream; nor does it place demands on lexical retrieval. Thus the two specific theories of interest make contrasting predictions about this task.

#### B. Phonological Decision

Participants were required to make a rhyme judgement on pairs of words represented by pictures. The pictures were easily nameable line drawings of single syllable words, mostly taken from the International Picture Naming Project (IPNP) database (https://crl.ucsd.edu/experiments/ipnp/index.html, Szekely et al., 2004). The pictures were arranged into 45 rhyming and 45 non-rhyming pairs (based on pairings devised by Bishop & Robson, 1989). Rhyming and non-rhyming pairs did not differ significantly on orthographic similarity (assessed using MatchCalc software, http://www.pc.rhul.ac.uk/staff/c.davis/Utilities/MatchCalc/). For each trial, a series of 6 picture pairs was presented, each for 3.33 seconds (totaling 20 seconds). For each pair, the participant decided whether the words represented by the pictures rhymed or not, and responded by button press.

This task involves implicit generation of lexical items and their phonology, but does not require access to conceptual meaning. Both the dorsal-ventral stream theory and lexical retrieval theory predict it should be strongly lateralized.

#### C. Semantic Decision

This task involved a semantic category judgement on objects represented in a pair of pictures.

The design of this task closely matched that of the phonological decision task. The pictures were mostly taken from the IPNP database, as described above. The stimuli were matched for word familiarity, orthographic neighbourhood, imageability, number of phonemes and frequency. Six picture pairs were presented, each for 3.33 seconds. For each pair, the participant decided whether the objects were from the same semantic category or not (e.g. both types of food) and responded by button press. For this task, it is necessary to access conceptual meaning, but generation of word names is not implicated. This, then, can be regarded as indexing the ventral stream. Both the dorsal-ventral stream theory and the lexical retrieval theory predict weak lateralization for this task.

#### D. Sentence Generation

This task required participants to generate spoken sentences in response to line drawings, following methods described by Mazoyer and colleagues (Mazoyer et al., 2014), but using pictures that were more culturally appropriate for UK participants.

For each trial, a black line drawing was displayed on a grey background for 3 seconds. This was followed by a fixation cross for 11 seconds, during which the participant was required to covertly generate a sentence. Participants were trained in advance to generate sentences beginning with a subject (e.g. “the boy”), followed by a description of the subject (“with marbles”), a verb (“plays”) and ending with a detail about the action (“on the floor”). A “REPORT” prompt was then presented for six seconds, and participants were required to say their sentence aloud.

This task implicates both dorsal and ventral streams, and so might be expected to show weaker lateralization than purely dorsal tasks. In contrast, the lexical retrieval theory predicts strong lateralization.

#### E. Sentence Comprehension

This task required participants to decide which of two pictures corresponded to a spoken sentence. Each trial comprised six picture pairs, each presented for 3.33 seconds, along with a spoken sentence that matched one of the two pictures. The sentences were spoken at a rapid pace and included some involving complex grammar with long-distance dependencies, such as ‘the shoe on the pencil is blue’, or ‘the cow that is brown is chasing the cat’. Participants indicated which of the two pictures matched the sentence by button press.

This task would appear to stress the ventral more than the dorsal stream, and so be relatively weakly lateralized. The task is hard to categorise in terms of lexical retrieval: it is necessary to hold word meanings in memory while working out the meaning, though overt word generation is not required.

#### F. Syntactic Decision

This task was designed to isolate syntactic processing with minimal involvement of semantics. This task uses ‘Jabberwocky’ stimuli, based on a study by Fedorenko and colleagues (Fedorenko, Hsieh, Nieto-Castañón, Whitfield-Gabrieli, & Kanwisher, 2010), where content words of sentences are replaced by plausible non-words. Half of the stimuli were ‘sentences’, where function words, word order and morphological cues were preserved to make the stimuli recognisable as syntactically-valid sentences (e.g. ‘The tarben yipped a lev near the kruss’). The other half had a pseudorandom word order and were not perceived as sentences (e.g. ‘Kivs his porla her tal ghep in with’).

Each trial contained three Jabberwocky stimuli of 8 words. Words were presented sequentially at the same time as an audio recording of the spoken word. As all spoken words were recorded separately, there were no prosodic cues to whether the stimulus is a ‘sentence’ or not. Each word was presented for 0.7 seconds, and the sequence was followed by a question mark for 1 second (making a total of 6.7 seconds for each Jabberwocky stimulus). The participant was required to respond by button press following the ‘?’ prompt.

In terms of the dorsal-ventral stream account, this task is predicted not to show lateralization, as it is a purely receptive task. This was the only task involving nonwords, and should not be lateralized according to a lexical retrieval account.

### Behavioural Analysis

For tasks A and D, the average number of words generated for each trial was calculated. For tasks B, C, E and F, percentage accuracy and average reaction time for correct trials (excluding trials where reaction time was greater than 2 standard deviations away from the mean) were calculated. The number of events where no response was received was also recorded for each task– these events were scored as incorrect.

### fTCD Analysis

Our analysis of fTCD data departed from the method we preregistered in three respects; sections describing the altered methods are shown in italics, with a description and explanation of the change shown in the section ‘Departures from pre-registered methods’.

The dependent measures derived from the fTCD analysis were the Laterality Indices (LI) from tasks A to F at sessions 1 and 2. fTCD uses ultrasound probes positioned bilaterally over the temporal windows to measure cerebral blood flow velocity (CBFV) in the left and right middle cerebral arteries (MCA). The probes emit ultrasound pulses and detect reflected ultrasound signal. The frequency of the reflected ultrasound signal depends on the speed of the blood moving in the MCA, due to Doppler shift. Hence the difference in frequency of the emitted and reflected ultrasound signals can be used to determine the speed of blood flow. The data is recorded as CBFV (cm/s) in the left and right hemispheres.

The fTCD data were analysed using a custom script in R Studio (RStudio Team, 2015). The script can be found on OSF (https://osf.io/tkpm2/). The CBFV data was first down-sampled from 100 Hz to 25 Hz by taking every 4^th^ datapoint. The data was segmented into epochs of 33 seconds, beginning 7 seconds before the presentation of the ‘CLEAR’ stimulus at the start of the trial (−7 seconds peri-stimulus time). Spiking or dropout datapoints were identified as being outside of the 0.0001 - 0.9999 quantiles of the CBFV data. If only a single artifact datapoint was identified within an epoch, it was replaced with the mean for that epoch. If more than one datapoint was identified, the epoch was rejected. The CBFV was then normalized (by dividing by the mean and multiplying by 100) such that the values for CBFV become independent to the angle of insonation and the diameter of the MCA. Heart cycle integration was used to normalize the data relative to rhythmic modulations in CBFV. *Each epoch was baseline corrected using the interval from −5 to 2 seconds peri-stimulus time*. Finally, artifacts were identified as values below 60% and above 140% of the mean normalised CBFV – any epochs containing such artifacts were rejected.

If a participant in one session had fewer than 12 acceptable epochs for any task (i.e. more than 3 of the 15 epochs were rejected), the data for that task were excluded. If a participant had more than one task excluded, all data for that participant were excluded.

The CBFV from left and right sensors was averaged over all epochs at each timepoint, and the mean difference (left minus right) within the period of interest was taken as the laterality index (LI). The period of interest for tasks B, C, E and F was from 6 to 23 seconds peri-stimulus time. For tasks A and D, the period of interest ended at 17 seconds to avoid activity related to overt speech production following the ‘REPORT’ prompt.

The LI value at each trial was also recorded, and used to calculate a standard error, which indicated how variable the lateralization was over trials. Outlier standard error values were identified using Hoaglin and Iglewicz’s procedure (Hoaglin & Iglewicz, 1987). The standard error values for every LI measurement (across all subjects, tasks and sessions; 360 values in total) were concatenated. The difference between the first and third quartiles of the data was calculated (Q3-Q1). In this dataset, outliers were defined as having standard error value more than 2.2 times this difference above the third quartile (Q3); e.g., the threshold limit = Q3 + 2.2*(Q3-Q1). Hence, if the LI value showed exceptionally high variability across trials, it was deemed to be unreliable and therefore omitted from the final analysis.

### Departures from pre-registered methods

#### 1. Baseline interval

The baseline interval was 2 seconds longer than that planned in the preregistered protocol (−5 to 0 seconds), i.e. extending into the ‘Clear mind’ period. As shown in Supplementary Materials (https://osf.io/g8mkv/), this baseline gives more stable estimates of LI than the original interval.

#### 2. Definition of laterality index

In our pre-registered protocol, we planned to use a peak-based method of measuring the Laterality Index (LI) developed by Deppe et al (Deppe, Knecht, Henningsen, & Ringelstein, 1997), which has been standard in fTCD studies of cerebral lateralization. This involves finding the absolute peak in the difference wave within the period of interest and averaging the value of the difference over a 2 second time window centered on this peak. The major limitation of this approach is that it creates a non-normal distribution of LI values, which contributed to poor model fit in our SEM analyses, which assume normality. The mean-based method that we report here gives LI values that are highly correlated with the traditional peak-based LI (Spearman r = 0.97), but with a normal distribution (see Supplementary Materials, https://osf.io/g8mkv/, for further details).

#### 3. Outlier detection

In our pre-registered document, there was an error in our description of this process; we mistakenly stated we would remove outliers based on LI scores, rather than the standard error of the LI scores. Removing LI outliers would not be sensible in the context of this study, where the focus is on individual differences: it would, for instance, lead us to exclude those with atypical right-sided language laterality, who are of particular interest for our hypothesis. Our goal in outlier removal was to exclude participants with noisy data, and the LI standard error is the appropriate measure to use to achieve this goal.

#### 4. SEM modelling

In addition to testing the models specified in the pre-registration document, we also tested model fit of the best-fitting model using a leave-one-out procedure, which allowed us to check whether the parameter estimates were unduly influenced by specific data-points. As described in Supplementary Materials (https://osf.io/g8mkv/), our decision to test further right-handers was prompted by discovering that there was undue influence from one left-hander, with the factor solution changing when her data were omitted. Accordingly, we present here additional analyses with 30 right-handers only, and with the full sample of 37 participants. We also computed the factor scores from the final model and plotted these to aid interpretation of the factor structure. The SEM bifactor model requires one variable to have fixed paths of 1 and 0 respectively to the two factors. The fit of the model does not depend on which measure is used for this purpose, but the specific path estimates will vary. Given that List Generation task was the only task with poor test-retest reliability, we present here results using Sentence Generation for the fixed paths. This follows recommendations that the strongest indicator for a specific factor should be used for the fixed paths (Lewis, 2017).

### Structural Equation Modelling

Structural Equation Modelling (SEM), as implemented in OpenMx (https://openmx.ssri.psu.edu/), was used to test our hypotheses. We distinguish between two sets of hypotheses: models of task effects, which concerned predictions about means, and models of person effects, which concerned covariances. As noted above, these are independent from one another. The models used to test each hypothesis are described below, and can be seen in Figure 4.

We will briefly describe this approach, as it not widely used in laterality research. The aim is to test how well a prespecified model fits an observed dataset. Typically SEM is used to model covariances, but it can also be used with means. Boxes denote observed variables, two-headed arrows show variances and covariances. A triangular symbol denotes a mean value, typically set to one, with the path from the box to the triangle corresponding to the mean value for that variable. Means can be set to be equivalent by giving their paths the same label. We use capital letters for paths to means. For instance, in the Population Bias model (Figure 4), all paths to the mean are set to be the same, whereas in the Task Effect model (Figure 4), the means differ from task to task, but within a task are the same from test session 1 to test session 2.

An oval symbol corresponds to a latent variable linking two observed variables: covariance between two observed variables is computed as the sum of the product of the paths to those variables that are linked by an oval. Paths to latent variables are shown as lower case letters. The difference between modeling of means and covariances can be appreciated by comparing the Task Effect model and the Person Effect model in Figure 4. These look similar, but the former depicts the situation where the means for a task are constant across sessions, but covariances are not considered. Thus even if means are stable, tasks may be unreliable in the sense that individual differences are just due to noise, and the rank order of LIs of individuals is unstable. In contrast, the Person Effect model takes into account covariances, and is a test of the reliability of the measures, assessing how far individuals are consistent in their LI across occasions.

We report goodness of fit for each model relative to a ‘saturated’ model where all variables are unconstrained, using the Comparative Fit Index (CFI): a high CFI indicates good model fit, and it is generally recommended that CFI needs to exceed .95 for the model to be regarded as a good fit to the data. We also report the Root Mean Square Error of Approximation (RMSEA), which is a measure of badness of fit, and should ideally be below .08 (Kline, 2011).

Comparison of model fit to determine the most appropriate model is achieved using likelihood ratio testing. Such comparisons are valid when we have nested models. For each hypothesis, we compare two nested models computing the difference in −2 log likelihoods, and evaluated in terms of the difference in degrees of freedom between the two models. The difference in log likelihoods follow a *χ*^2^distribution, so a *χ*^2^ test can be used to evaluate whether there is a statistical difference between the models. If a significant difference is found, then one model will be a better fit to the data.

In general, when comparing a model against another more complex model, good model fit corresponds to a non-significant p-value, which indicates that the more parsimonious model fits as well as the more complex model, despite fewer degrees of freedom. Models that estimate many parameters (and so have fewer degrees of freedom) will tend to fit the data better, and so relative fit of models is considered using indices that take this into account. Several indices that penalize the likelihood ratio test are available, for example, Akaike’s Information Criterion (AIC) or Bayesian Information Criterion (BIC). Both these indices provide a value for each nested model and the lowest value among all the models is the preferred model.

#### step 1: Testing Stability of LI Values

We began with a Fully Saturated model that modeled means and variances as totally independent, as shown in Figure 4 (top left). No correlations between LI values were modelled at this stage: the triangular symbol denotes that the paths reflect the mean for each observed variable. As an initial sanity check, we computed a Task Effect model where the LI value means and variances for each task (A-F) were fixed to be the same at each testing session (i.e. the means and variances for A1 = A2, B1 = B2, etc.). We predicted that the latter model would not deteriorate compared to the Fully Saturated model, indicating that we would not need to specify separate means for different test occasions.

#### step 2: Testing Models of Means

Our first hypothesis proposed that a significant task effect on LI value would be observed; i.e., that the mean LI values would vary between the six different tasks (tasks A-F). This was assessed by comparing the two models shown in row 2 of Figure 4: the Population Bias model and the Task Effect model.

The Task Effect model was then used as a baseline comparison model to test two more specific sub-hypotheses regarding which tasks would show the strongest lateralisation. In each case we divided tasks into three subsets, and fixed the means and variances for the tasks within each subset to be the same. We adopted this approach to test the Dorsal Stream hypothesis and the Lexical Retrieval hypothesis.

#### step 3: Testing Models of Covariances

Two models of covariance were compared (Figure 4, bottom). First, a person effect model was computed where covariance was predicted by a single factor, i.e. was similar across all language tasks. This was compared with a person by task effect model, with two covariance factors. The Person Effect (single factor) model is nested within the Task x Person Effect (bifactor) model, and so their relative fit can be assessed by subtraction of negative log likelihoods.

All analyses were conducted in R (R Core Team. & R Development Core Team, 2013). Data and analysis script are available on Open Science Framework.

**Figure 4:**
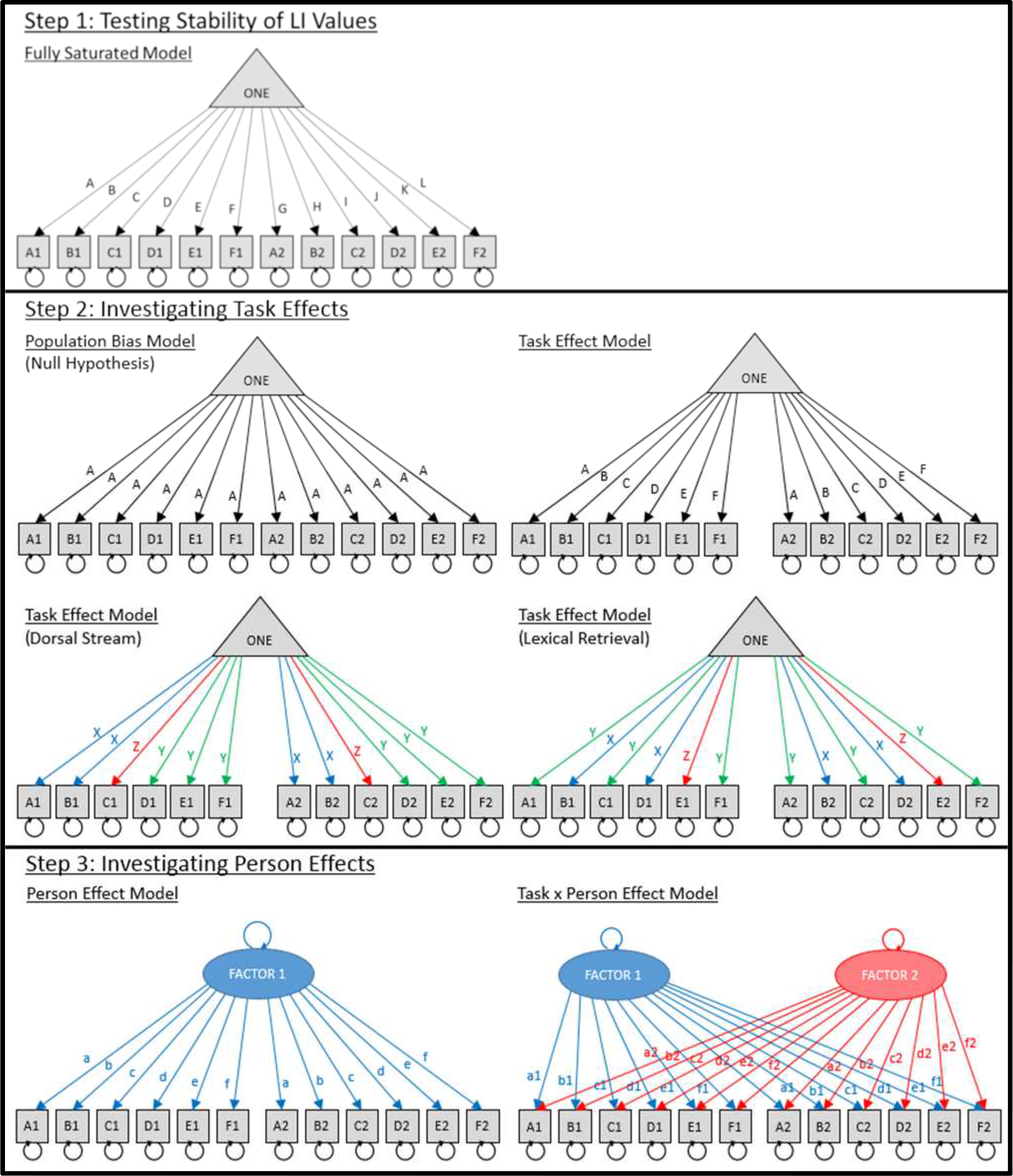
Step 1 (top): Simple model of means and variances. In the ‘Fully Saturated’ model the means for all tasks could vary independently (tasks A-F, tested at sessions 1 and 2). This was compared to the ‘Task Effect’ model, where the means for each task were fixed to be the same for each session. The triangle symbol denotes that this is a model of means: covariances between values are not included in the model.
Step 2 (middle): To test hypotheses relating to the LI means, the ‘Population Bias’ model (with means for all tasks set to be the same) was compared to the ‘Task Effect’ model (where means varied by task). Furthermore, to test the ‘Dorsal Steam’ hypothesis, a model with means for subsets of dorsal (A, B), ventral (C) and mixed tasks (D, E, F) were fixed (labelled as X, Z and Y). For the ‘Lexical Retrieval’ hypothesis, a model with means for subsets of tasks with lexical retrieval (B, D) and tasks without (A, C, F) were fixed (labelled as X and Y respectively). Step 3 (bottom): The oval symbol denotes a common factor that determines the covariance between observed variables. To test the hypothesis relating to LI covariances, a single factor ‘Person Effect’ model, was compared to a two factor ‘Task x Person Effect’ model. To achieve model identification, one of the paths from Factor 1 to a task had to be fixed to 1, and the path from Factor 2 to that task was fixed to zero. *In our preregistration this fixed path was planned to be task A, but due to the low reliability of that task, it was changed in the final analysis to be task D.* The covariance between Factor 1 and Factor 2 was also set to zero. Note that the means were also modelled as shown in the task effect model, but this was omitted from the model diagrams here for simplicity.

## Results

All data are available on OSF (https://osf.io/s9kx6/). Results from the pre-registered analysis protocol (i.e., using the first 30 participants only) are shown in Supplementary Materials (https://osf.io/g8mkv/). As noted above, the factor solution from this sample was unstable and unduly influenced by one left-hander. We report here the results based on the final sample of 30 right-handers and 7 left-handers, which gives a stable solution, and we include exploratory analyses relating the findings to handedness. The LI values reported here are based on the mean difference between left and right CBFV, as this gives normally distributed variables, but the results are highly similar when the non-normal peak-based LIs are used instead. The analysis script provided on OSF (https://osf.io/q8zka/) facilitates comparisons between different analytic pathways.

### Behavioural results

We did not have specific predictions for the behavioural results, but present them here for completeness. For List Generation (A) and Sentence Generation (D), the number of words spoken per trial was recorded. The number of words spoken in both tasks and sessions were very similar: for task A, session 1, mean = 9.5, SD = 0.42, session 2, mean = 9.6, SD = 0.29; for task D, session 1, mean = 9.2, SD = 1.21, session 2, mean = 9.4, SD = 1.24. A repeated measures ANOVA showed no significant effects of task (F(1,36) = 1.22, *p* = 0.278) on the number of words spoken, but there was a significant effect of session (F(1,36) = 5.73, *p* = 0.022). Trials where participants failed to respond, or responded too early were excluded from analysis: these constituted less than 0.1% of trials.

For decision making tasks (B, C, E and F), the accuracy and RT of each response, and the number of omitted responses, were recorded (Table 1). Note that for task F participants were required to wait until the end of the word sequence before responding, and had only a second to respond; this accounts for the fast reaction times and relatively high number of omitted responses in task F.

The Phonological Decision and Sentence Comprehension tasks (tasks B and E) showed evidence of practice effects, as both accuracy and reaction times improved, and the number of omitted responses fell from Session 1 to Session 2.

**Table 1.** Behavioural data for tasks B, C, E and F. The table shows mean percentage accuracy and reaction times (with SD), and results of t-tests comparing Session 1 with Session 2 for each measure. The number of omitted responses is reported as a percentage of all events. B = Phonological Decision; C = Semantic Decision; E = Sentence Comprehension; F = Syntactic Decision.

| Measure | Session | Task B | Task C | Task E | Task F |
| --- | --- | --- | --- | --- | --- |
| Accuracy (%) | 1 | 91.3 (5.55) | 95.9 (3.08) | 92.5 (4.81) | 89.6 (8.31) |
|  | 2 | 93.3 (4.28) | 95.0 (3.06) | 94.2 (3.79) | 89.4 (8.28) |
|  | 1 vs 2 | t=-3.27, p=.002 | t=1.61, p=.115 | t=-2.70, p=.011 | t=-0.07, p=.944 |
| Reaction times (s) | 1 | 1.66 (0.22) | 1.14 (0.2) | 2.17 (0.12) | 0.33 (0.08) |
|  | 2 | 1.49 (0.21) | 1.05 (0.2) | 2.11 (0.15) | 0.33 (0.07) |
|  | 1 vs 2 | t=8.73, p<.001 | t=4.77, p<.001 | t=3.27, p=.002 | t=0.64, p=.528 |
| Omitted responses (%) | 1 | 2.34 | 0.84 | 2.79 | 4.20 |
|  | 2 | 0.78 | 0.60 | 1.62 | 4.44 |

### Lateralisation results

Three outlier LI values were excluded where the standard error across trials was above the upper cut-off. Six LI values were excluded because a subject had less than twelve useable trials for a given task in a given session. The remaining data for these participants were retained in the analysis. Excluded datapoints are shown as red dots in Figure 5.

Figure 5 shows the distribution of LIs as a pirate plot (Phillips, 2017). Task D (Sentence Generation) showed the strongest left lateralisation. Shapiro-Wilks normality tests showed that LI values for all 12 conditions were normally distributed. One sample t-tests (testing for mean > 0) showed that all conditions were significantly left lateralised, except task F (Syntactic Decision; Session 1: t (33) = 0.77, p = 0.224; Session 2: t (36) = 0.33, p = 0.373).

**Figure 5.**
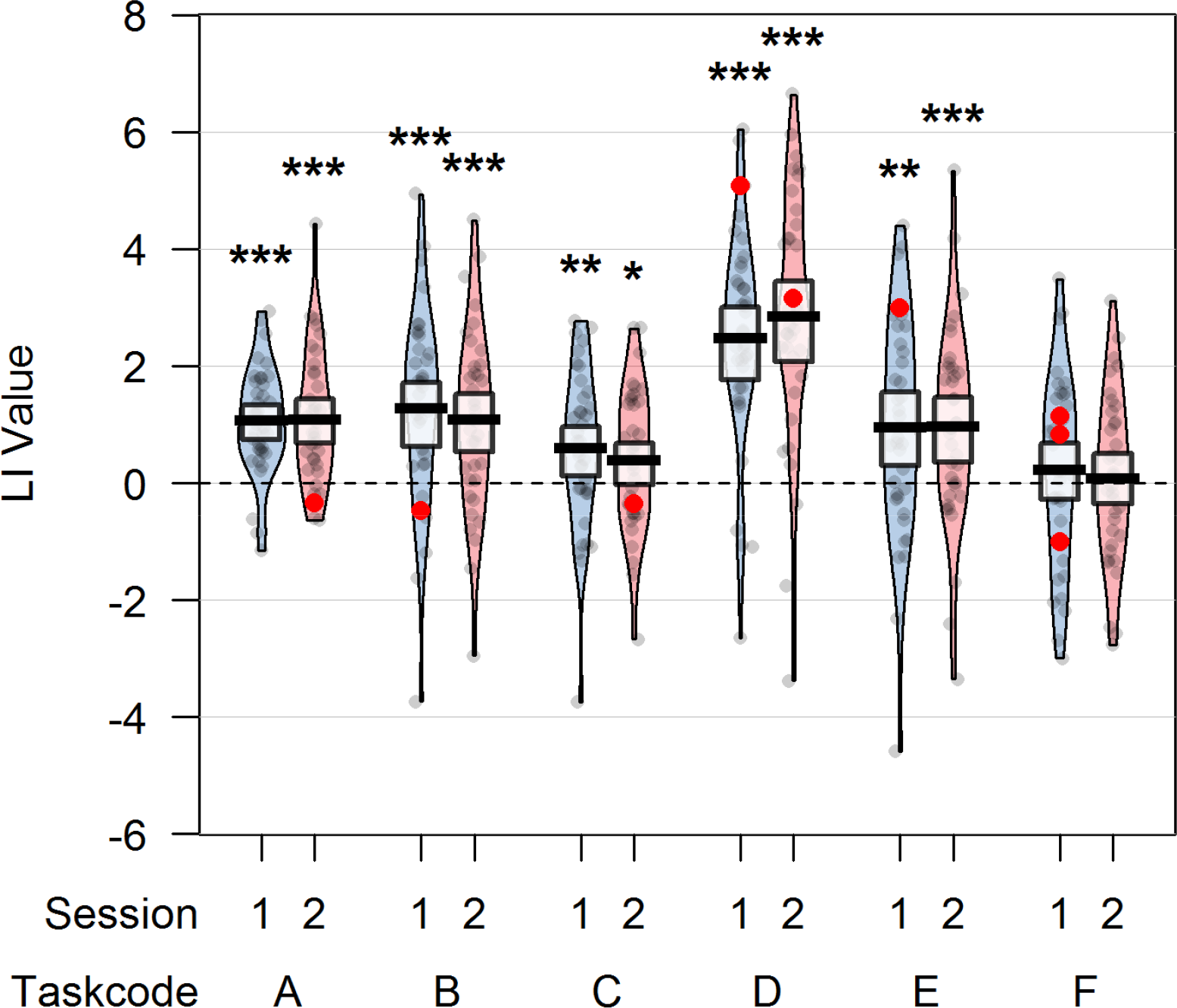
Pirate plot of LI values for all tasks (A-F) and sessions (blue = Session1, pink = Session2). Excluded data-points are shown in red. Asterisks show results of Wilcoxon tests comparing the LI values of the group (omitting excluded data-points) to zero (* *p*<.05; ** p<.01; *** p<.001).

Figure 6 shows a correlation matrix of LI values for all tasks and sessions. Test-retest correlations varied between tasks. Task A (List Generation) had poor test-retest reliability (Pearson’s r = 0.13), and low correlations with other tasks. Test-retest reliability for other tasks ranged from r = 0.57 to 0.84. Tasks B, C, D and E were strongly intercorrelated. Task F (Syntactic Decision) had high test-retest reliability (r = 0.76) but relatively low correlations with other tasks.

**Figure 6.**
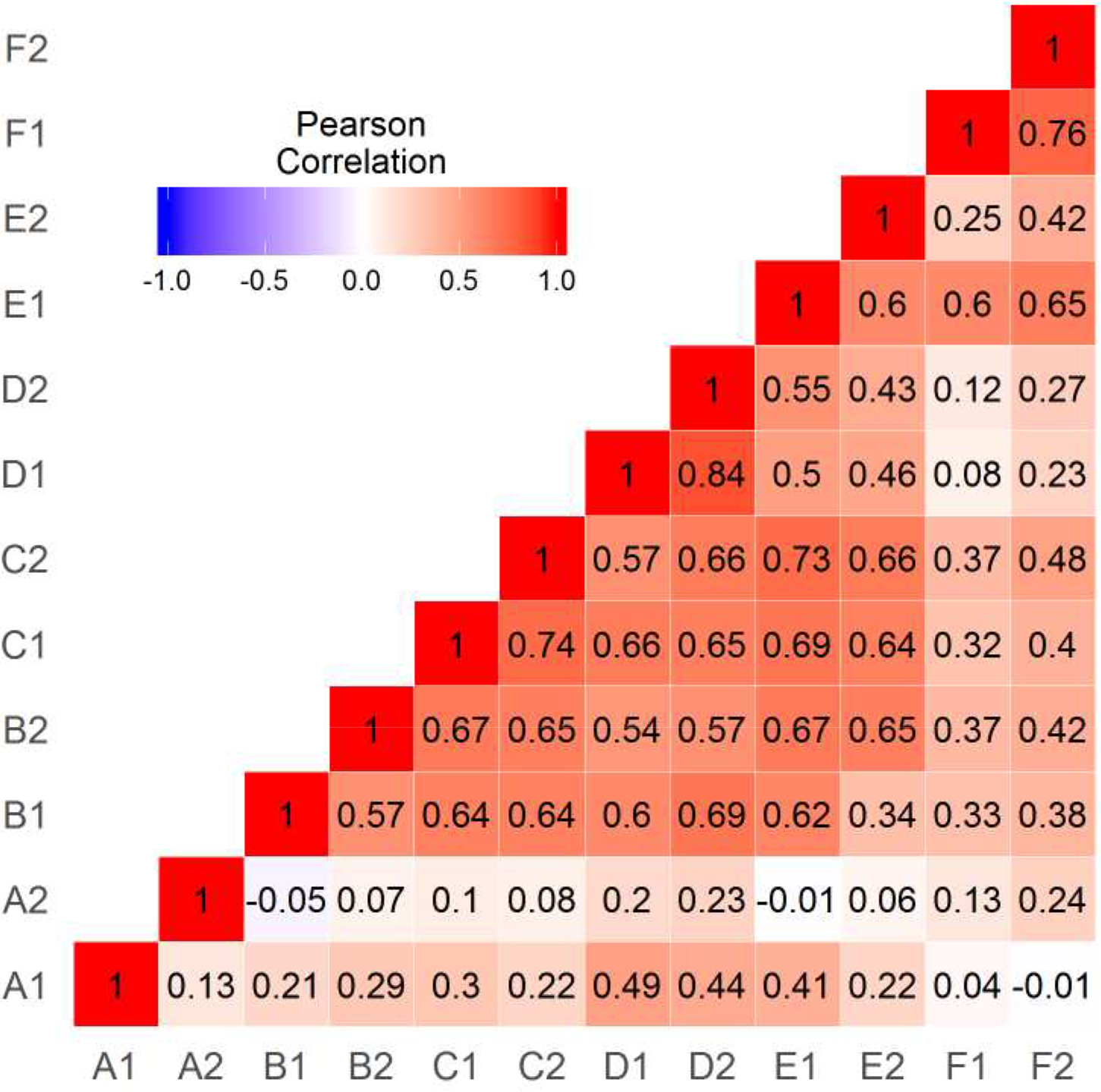
Correlation matrix for LIs from the six language tasks given on two occasions.

### Structural Equation Modelling

The LI data were entered into the SEM analysis to test hypotheses about the group mean LI values and covariances in LI values across subjects. Table 2 summarises the SEM results.

#### step 1: Testing Stability of LI Values

As shown in Table 2, the fit of all the means-only models was very poor. This is to be expected, as these models ignore covariances, and, as indicated in Figure 6, there are substantial correlations both between and within tasks. Our interest at this point, however, is in the relative fit of different models of means, rather than overall model fit. The Fully Saturated model (with free means and variances) was compared to the Task Effect model, which fixed the means and variances for each task to be stable over sessions (i.e. A1 = A2, B1 = B2, etc.). The Task Effect model fit did not deteriorate significantly from that of the Fully Saturated model, supporting the hypothesis that LI means for each task were stable across sessions.

**Table 2.** Model fit statistics from structural equation models and model comparisons. −2LogL = −2 log likelihoods; df = degrees of freedom; BIC = Bayesian Information Criterion;CFI= Comparative Fit Index; RMSEA = Root Mean Square Error of Approximation.

| Model | Description | -2LogL | df | BIC | CFI | RMSEA | Chi Square test<br>Compared to | p |
| --- | --- | --- | --- | --- | --- | --- | --- | --- |
| Fully Saturated Model | Free means and variances | 1574.4 | 411 | 90.4 | NA | NA | - | NA |
| Task Effect Model | Stable means and variances | 1580.8 | 423 | 53.4 | 0.022 | 0.292 | Fully Saturated Model | 0.896 |
| Population Bias Model | Equal means and variances | 1715.5 | 433 | 151.9 | -0.474 | 0.337 | Task Effect Model | <0.001 |
| Dorsal Stream Model | Means for tasks AB > DEF > C | 1664.8 | 429 | 115.7 | -0.288 | 0.323 | Task Effect Model | <0.001 |
| Lexical Retrieval Model | Means for tasks BD > ACF | 1631.6 | 429 | 82.5 | -0.156 | 0.306 | Task Effect Model | <0.001 |
| Person Effect Model | Covariances have one factor structure | 1378.4 | 417 | -127.4 | 0.805 | 0.136 | Task Effect Model | <0.001 |
| Task x Person Effect Model | Covariances have bifactor structure | 1337.8 | 412 | -149.9 | 0.947 | 0.073 | Person Effect Model | <0.001 |

#### step 2: Testing Models of Means

To demonstrate whether LI means differed between tasks, the Task Effect model (with different means for each task) was compared to the Population Bias model (with means fixed to be the same for all tasks). This may be seen as a null hypothesis that treats all tasks as equivalent measures of laterality. The Population Bias model gave significantly worse fit (see Table 2), supporting the hypothesis that LI means differed between tasks.

Two further models were compared to the Task Effect model. The Dorsal Stream model categorised the language tasks according to the involvement of the dorsal or ventral stream. Tasks A and B were categorised as involving strong dorsal stream activity, task C as strong ventral stream activity, and tasks D, E and F as intermediate (hence, means for AB > DEF > C). This model gave significantly poorer fit than the Task Effect model – as is evident from Figure 5, which shows relatively weak lateralisation for tasks A and B compared to task D. The Lexical Retrieval model did not fare any better. This categorised tasks B and D as involving strong lexical retrieval, whereas tasks A, C and F did not involve lexical retrieval, and task E was difficult to classify and so was considered as independent of the other measures (BD > ACF). Again, this model gave a worse fit than the Task Effect model, indicating that, while laterality varied between tasks, it did not fit the either of the predicted patterns. Note, however, that the pre-registered tests specified for both theories have some limitations, as discussed further below.

#### step 3: Testing Models of Covariances

At Step 3 we tested whether the covariances between tasks had a single factor structure (Person Effect model) or a bifactor structure (Task by Person Effect model). Not surprisingly, given the strong correlations in Figure 6, both within and across tasks, the Person Effect model gave substantially better fit than the Task Effect model (see Table 2); nevertheless, the overall fit of this model was poor. The Task by Person Effect model gave a significantly improved fit. A plot of the two factors is shown in Figure 7: note that, although the model fit is not affected by task selection, the factor scores depend on which task has fixed paths to the factors. The paths for the case when Sentence Generation is fixed are shown in Table 3. It can be seen that List Generation has only a weak loading on Factor 1, whereas Phonological Decision, Semantic Decision and Sentence Comprehension have moderate loadings on both factors. Syntactic Decision has a strong loading on Factor 2 but does not load on Factor 1, reflecting the weak correlation of this task with Sentence Generation.

**Table 3.** Path weightings (and 95% confidence intervals) from each latent factor (Factor 1 and Factor 2) to each task (A to F) from the winning bifactor model.

| Task | Factor 1 |  | Factor 2 |  |
| --- | --- | --- | --- | --- |
|  | Path | 95% CI | Path | 95% CI |
| A: List Generation | 0.18 | 0.05 to 0.31 | -0.02 | -0.27 to 0.24 |
| B: Phonological Decision | 0.61 | 0.40 to 0.81 | 0.55 | 0.21 to 0.89 |
| C: Semantic Decision | 0.53 | 0.36 to 0.69 | 0.52 | 0.23 to 0.81 |
| D: Sentence Generation | 1.00 | Fixed | 0.00 | Fixed |
| E: Sentence Comprehension | 0.56 | 0.30 to 0.82 | 0.95 | 0.54 to 1.37 |
| F: Syntactic Decision | 0.13 | -0.13 to 0.40 | 1.16 | 0.75 to 1.56 |

In our original analysis with 30 participants, a similar factor structure was observed, but there was a concern that this depended solely on a single left-handed participant (see Supplementary Material, https://osf.io/g8mkv/). With the larger sample of 37 participants, the bifactor (Task by Person Effect) model was superior in all runs of a leave-one-out analysis. The bifactor model was also the best-fitting model when only the 30 right-handers were included in the analysis. Nevertheless, it is clear from Figure 7 that the two factors were highly intercorrelated, and the impression is that the bifactor solution is heavily affected by some influential cases. Cook’s distance identified four bivariate outliers, marked with circles in Figure 7: all four outliers were left-handers. When the analysis was re-run omitting these cases, the single factor model gave a better model fit when all N=33 subjects were included (single factor BIC=−142.7, bifactor BIC=−138.6), and in all but one run of the leave-one-out analysis.

We can conclude from this analysis that, although univariate normality was satisfactory, our data did not meet conditions of multivariate normality; this leads to the conclusion that the sample is not homogeneous, but contains a mixture of laterality patterns. We discuss the implications of this finding below.

**Figure 7.**
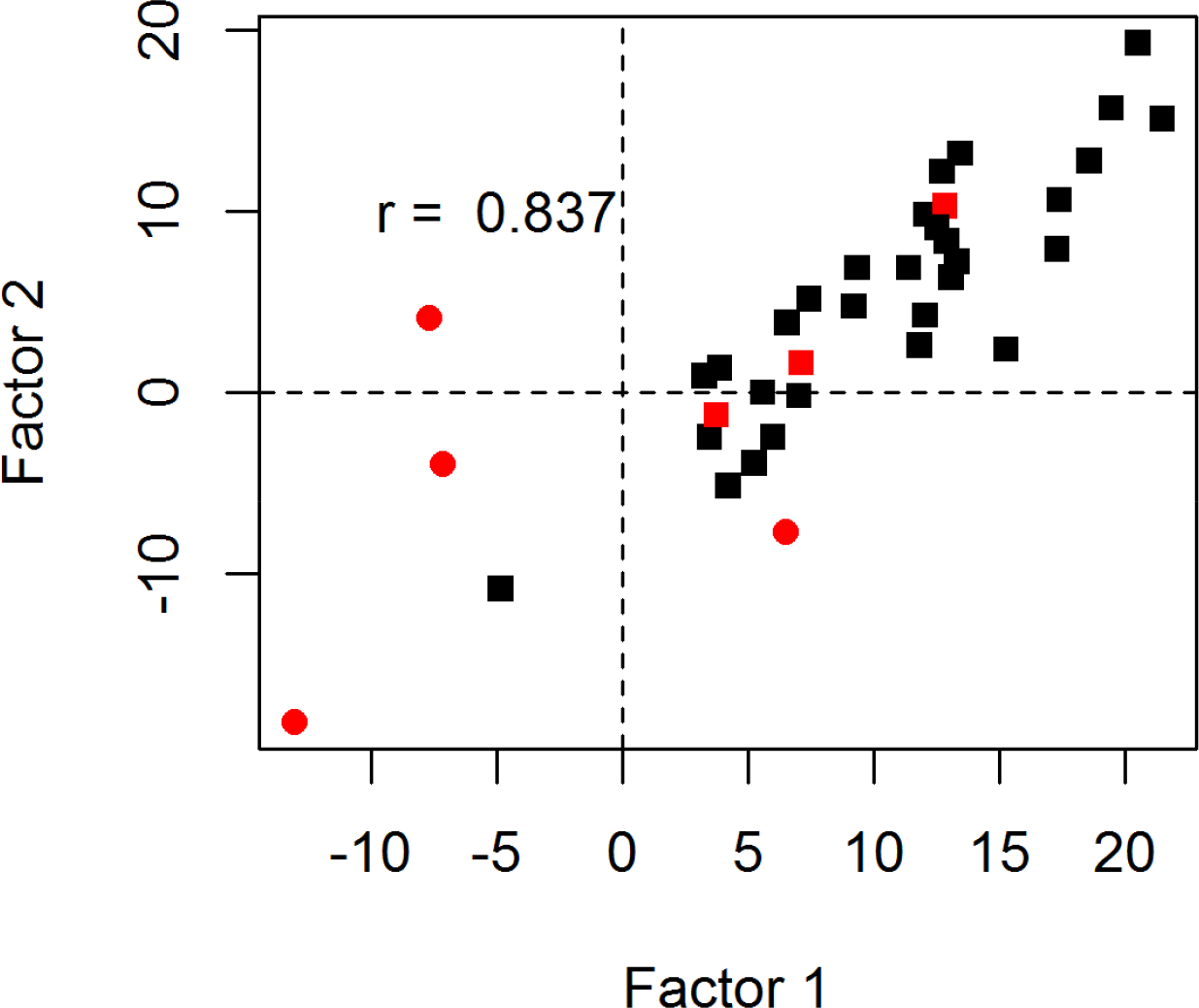
Correlation between two factors from the bifactor (Task by Person Effect) model, with left-handers shown in red, and bivariate outliers as circles.

## Discussion

The question of whether cerebral lateralisation is a unitary function may be interpreted at two levels: at the population level, we may ask whether all language tasks show a similar degree of lateralisation, and at the individual level, whether people show consistent differences in laterality profiles across tasks.

Although we used formal modelling to address these questions, a good insight into the answers can be obtained by viewing figures 5 and 6. Figure 5 shows clear differences from task to task in strength of cerebral lateralisation, whereas Figure 6 shows moderate-to-good test-retest reliability for all but one task, coupled with significant cross-task correlations.

The SEM analyses allowed us to explore these patterns further. Regarding means, as expected, a null hypothesis of no difference between tasks could be convincingly rejected. However, the specific patterns that we predicted should be seen on the basis of two existing models – the Dorsal Stream model and the Lexical Retrieval model – did not give a good fit. It could be argued that the data are, in fact, consistent with the Dorsal Stream model, insofar as the three tasks that involved implicit or explicit generation of speech – List Generation, Phonological Decision and Sentence Generation – were the ones that showed the strongest lateralisation (see Figure 5). The poor fit of the Dorsal Stream model was in part due to the fact that Sentence Generation was judged to implicate both streams, and was not therefore predicted to be as strongly lateralised as tasks with weaker semantic demands. However, it clearly makes demands on the phonological-articulatory system, and with hindsight it could be argued that in terms of articulatory complexity it was more demanding than the other tasks. A key question is whether blood flow measured using fTCD reflects the average of activity in a lateralised dorsal stream and a bilateral ventral stream, or whether the absolute dorsal stream activity is the main factor affecting the LI. In future we plan studies to address this question using fMRI.

More generally, based on the pattern of results observed in this study, it appears that whole-hemisphere lateralisation as measured by fTCD is driven most strongly by generation of meaningful, connected speech (e.g. Sentence Generation). Lateralization for this task was stronger than for automatic, non-propositional speech (List Generation) or implicit sub-vocalisation (Phonological Decision). By contrast, lateralisation was non-significant for the Syntactic Decision task.

We would, however, emphasise the need for caution in treating any one task as an indicator of a particular language function: it is evident that even minor modifications to task demands may affect laterality, particularly when sample size is relatively small. For instance, in a related study with a different sample of people, we recently found that List Generation was not lateralised (Woodhead, Rutherford, & Bishop, 2018). In that study we interleaved a simple number generation (counting) task with trials of Sentence Generation, whereas in the current study, List Generation was administered in a separate block, with the type of list (numbers, days of the week, months of the year) varied to engage the participants’ attention throughout the block. Although the counting task used by Woodhead et al (2018) was not significantly lateralised, it had good split-half reliability and was significantly correlated with Sentence Generation, whereas the List Generation task used in the current study was the only task to show poor test-retest reliability and relatively weak correlations with other tasks. Furthermore, our Semantic Decision task was designed to tap into similar semantic processes as the Pyramids and Palm Trees test (Howard & Patterson, 1992), but resulted in weaker LIs than seen in a study by Bruckert (Bruckert, 2016) using the Pyramids and Palm Trees task. It could be that the two-alternative forced choice task used in that study was more demanding than our match/no-match decision, but this kind of difference cautions us about relying on a single test to indicate a type of linguistic processing.

One convincing point to emerge from the analysis of mean data is that most language tasks (B, C, D, E and F) showed stable lateralisation measured in different sessions, but they differed in terms of the strength of left-lateralisation.

We turn next to the findings concerning covariances. It has been argued that fTCD is not useful for studying cerebral lateralisation because it is unreliable (Cai et al., 2013), but our data support those of Stroobant and Vingerhoets (Stroobant & Vingerhoets, 2001) in demonstrating that there is significant individual variation in language laterality between people that cannot just be attributed to noise. Furthermore, by moving from a definition of laterality based on a peak in the L-R difference wave to a definition based on mean L-R difference within a period of interest, we avoid the problem that can arise when laterality is forced into a non-normal distribution (see also Woodhead et al., 2018). As shown in Figure 5 and our tests of normality, when mean L-R difference is used, the distribution of LI values is normal.

The SEM also tested whether a single factor could explain individual differences in language lateralisation. At first glance, the results suggested this was not the case: the bifactor (Task by Person Effect) model showed superior fit over a single factor (Person Effect) model. This was the conclusion suggested by our initial pre-registered analysis, based just on a sample of 30 individuals. A leave-one-out analysis, however, made us cautious about accepting that result at face value, because the factor structure changed when a single left-hander with strongly complementary laterality on two tasks was excluded. For this reason we collected more data, adding seven right-handers to the sample. With this larger sample, we again found superiority for a bifactor solution, regardless of whether we included only right-handers or the full sample including left-handers. Yet there remained misgivings about the generalisability of the result, not least because the two factors were highly correlated (Pearson’s r = 0.84). A scatterplot of the two factors revealed a number of bivariate outliers and, as with our initial analysis, the pattern of results relied on which participants were included. Of course, it is not surprising that removing participants with the strongest dissociation between factors changes the factor structure: the point we wish to make is not that the results can alter in this way, but rather that the pattern of our SEM findings appears driven by heterogeneity within the sample, reflected in the presence of bivariate outliers.

The answer to the question of whether laterality is a unitary function is that, clearly, there are some individuals in whom laterality is different for different aspects of language. It is not, however, the case that there are two factors that act independently in the general population. Rather, the majority of people appear to have language laterality driven by a single process affecting all types of task, with a minority showing fractionation of language asymmetry.

The pattern of results is consistent with accounts of laterality that postulate qualitative rather than just quantitative differences between individuals. Theoretical accounts have mostly focused on a single dimension, arguing for laterality subgroups on the basis of non-normal distributions of scores (e.g. Mazoyer et al., 2014). Our results suggest that atypical laterality may be easier to identify when more than one language measure is considered, as detection of bivariate outliers can be effective with smaller samples than those required for detecting mixtures of distributions.

An association between atypical laterality and left-handedness has been established for many years, ever since early observations were made of superior recovery from aphasia after gun-shot wounds in left-handers (Subirana, 1958). However, most of the emphasis has been on atypical laterality in the sense of having language mediated by the right hemisphere. Although the number of left-handers in our sample is too small for numeric analysis, the fact that three of the four bivariate outliers were left-handers is a striking departure from chance (Fisher exact probability = 0.016) and compatible with the idea that language lateralisation is more likely to be multifactorial in left-handers than right-handers.

Further studies are needed to establish the key characteristics of tasks that index the two factors seen in some people, but we offer here some speculations. The main contributor to the second factor was the Syntactic Decision task, which differed from the other tasks in several regards. It used unfamiliar, nonword stimuli, and required the listener to identify syntactic errors. It was one of two receptive language tasks that involved processing of auditory language: the other was sentence comprehension, which had moderately strong loadings on the second factor. Perhaps the most surprising finding from this study is the fact that the one task that loaded on to the second factor (Syntactic Decision) was not lateralised, yet showed high test-retest reliability (R=0.67). We had anticipated that a lack of lateralisation on a task might be a consequence of noisy data giving poor test reliability – or alternatively a lack of individual variation if both hemispheres contributed equally in most people. Our data suggest that individuals do vary in the hemisphere used when doing the syntactic judgement task, and that this bias is reliable, but that it is not systematic across the population. This is perhaps the best evidence to date that strength as well as direction of lateralisation for a task is a stable trait.

### Limitations

As noted above, the principal limitation of fTCD is that it does not allow one to localise lateralised activity within a hemisphere. In future work, we plan to extend this line of investigation to consider whether similar patterns of lateralisation can be seen using comparable tasks with fMRI. The benefit of fTCD is that it is relatively inexpensive and quick to administer, and so enables us to gather data that can be used as a basis for developing a more hypothesis-driven approach that can then be extended and validated with fMRI.

A further limitation is that we lacked statistical power or range of measures that would be needed to evaluate more complex models. The bifactor model that gave the best fit in our study must be interpreted with caution. It will need to be replicated in larger samples and shown to generalise to new tasks - it remains a possibility that using a different set of tasks would reveal different or further fractionation of language lateralisation. Furthermore, although we have shown a bifactor model is a better fit than a single factor model, it is possible that more than two factors are needed to explain the full range of patterns of language lateralisation.

### Summary

In summary, these results indicate that there are meaningful differences in language lateralisation between tasks, and meaningful individual variability in lateralisation that is not simply due to measurement error. Even when a language-related task is not left-lateralised, there are stable individual differences in the contribution of the two hemispheres. Structural equation modelling of individual variability indicated that although a two-factor model gave a better fit than a single factor model, the effect was driven by a small subset of participants with discrepant laterality, and a single factor could account for variation in the majority of participants. Overall, our findings suggest there are qualitative as well as quantitative differences between people in laterality across tasks, and that consideration of asymmetry profiles on several tasks together can help identify cases of atypical laterality.

## Competing Interests

None to declare.

